# Multiple timescales account for adaptive responses across sensory cortices

**DOI:** 10.1101/700062

**Authors:** Kenneth W. Latimer, Dylan Barbera, Michael Sokoletsky, Bshara Awwad, Yonaton Katz, Israel Nelken, Ilan Lampl, Adrienne Fairhall, Nicholas J. Priebe

**Keywords:** adaptation, auditory cortex, somatosensory cortex, visual cortex, neural coding, linear-nonlinear model

## Abstract

Sensory systems encounter remarkably diverse stimuli in the external environment. Natural stimuli exhibit timescales and amplitudes of variation that span a wide range. Mechanisms of adaptation, ubiquitous feature of sensory systems, allow for the accommodation of this range of scales. Are there common rules of adaptation across different sensory modalities? We measured the membrane potential responses of individual neurons in the visual, somatosensory and auditory cortices to discrete, punctate stimuli delivered at a wide range of fixed and nonfixed frequencies. We find that the adaptive profile of the response is largely preserved across these three areas, exhibiting attenuation and responses to the cessation of stimulation which are signatures of response to changes in stimulus statistics. We demonstrate that these adaptive responses can emerge from a simple model based on the integration of fixed filters operating over multiple time scales.

## Introduction

Natural stimuli encompass, across all sensory modalities, a very wide range of amplitudes, and share structure at many spatial and temporal scales (Simoncelli & Olshausen, 2001, Fairhall 2014). We suggest that as a consequence, all sensory systems are subject to multiple mechanisms of adaptation that modulate their response properties over a variety of timescales ranging from milliseconds to hundreds of seconds. These adaptive modulations are driven by a number of factors such as the history of variations in input, statistical properties of the stimulus and the overall activity of the system. As a result, an encoding model developed for a given set of stimulus dynamics often fails to predict responses when the statistics of the stimulus change (Ozuysal & Baccus, 2012; Heitman et al., 2016; McIntosh et al., 2016).

While the statistical details differ across sensory modalities, natural stimuli all exhibit temporal fluctuations that are distributed across timescales. One may therefore expect the adaptive properties of sensory systems to be tuned to address these temporal fluctuations (Fairhall, 2014). Circuitry and cell types are similar across cortical fields as well, raising the question whether these similarities lead to common adaptive properties across different sensory areas. With this in mind, we explore here temporal properties of adaptation, documenting the dynamics of response sensitivity across sensory modalities.

To examine sensory adaptation across the neocortex, we investigated three sensory modalities - somatosensory, visual, and auditory. A major challenge in previous work has been that while the descriptions of adaptation for each of these systems are extensive, they are difficult to compare due to disparate experimental paradigms. Our goal here was to characterize neuronal responses across the sensory cortex to a common stimulus set, allowing us to move beyond modality-specific descriptions. To this end, we studied sequences of discrete punctate pulses (in the form of monitor flashes, auditory clicks and transient whisker deflections) delivered both at fixed frequencies and in Poisson-noise sequences. For each stimulus modality, we performed in vivo whole cell recordings in the respective cortical area (V1, S1, and A1).

We uncover a set of underlying fixed sensory filters that allows sensory neurons to adjust their sensitivity to temporally varying stimulus conditions. While the dynamics of neuronal responses appears to depend on stimulus conditions, we demonstrate that a common model, composed of filters with multiple timescales, can account for their emergent behavior. We found that major adaptive features of the membrane potential responses to fixed frequency stimuli are generally conserved across different cortical sensory modalities, though they differed in the degree to which they are expressed. These include a shift in the complexity of individual responses with the rate of stimulation, a reduction in response amplitude with the rate of stimulation, and a termination response at the end of high frequency stimulation. All of these components of the adaptive responses obtained from sensory neurons can be accounted for by a fixed, time-invariant model, indicating that adaptive processes for different sets of stimuli need not be the result of a change in the state of the network, but instead simply reflect the integration of multiple timescales of static sensory filters.

## Methods

### Physiology

Physiological procedures for mouse recordings are based on those previously described (Cang et al., 2008; Scholl et al., 2013). All of our experiments were conducted using adult C57BL/6 mice (P28-P90 to avoid the hearing loss that develops in older mice of this strain). Mice used in V1 experiments were aged P35 and older to avoid the visual critical period. Mice were anesthetized with 1,000 mg/kg urethane and 10 mg/kg Chlorprothixene via intraperitoneal injection. A further intraperitoneal injection of 20 mg/kg dexamethasone was administered to prevent brain edema. During the course of the experiment body temperature was monitored and maintained at 37°C. A tracheotomy was performed and the head was secured using custom-made head holders. A craniotomy and durotomy were performed over the appropriate area of sensory cortex. A1, S1 or V1 were located using standard techniques. Mouse eyes were kept moist with artificial tears or a thin layer of silicone oil. The cortical surface was kept moist with saline or 4% agarose in normal saline. All animal procedures were approved by the University of Texas at Austin Institutional Animal Care and Use Committee and by Animal Care and Use Committees at Hebrew University and the Weizmann Institute. Hebrew University is an AAALAC approved institution.

After the identification of the relevant area of sensory cortex, we performed in-vivo whole cell recordings using the blind patch method. A silver-silver chloride wire was inserted into muscle near the base of the skull and used as a reference electrode. For V1 and A1 recordings pipettes (5-10 MΩ) were pulled from 1.2 mm outer diameter, 0.7 mm inner diameter KG-33 borosilicate glass capillaries (King Precision Glass) on a P-2000 micropipette puller (Sutter Instruments). Pipettes were filled with (in mM) 135 K-gluconate, 4 NaCl, 0.5 EGTA, 2 MgATP, 10 phosphocreatine disodium, and 10 HEPES, pH adjusted to 7.3 with KOH(Sigma-Aldrich). For S1 recordings pipettes (1.5 mm outer diameter, 0.86 inner diameter, BF150-86-10, Sutter instruments) were pulled on a PC-10 vertical puller (Narashige) and were filled with (in mM):136 K-gluconate, 10 KCl, 5 NaCl, 10 HEPES, 1 MgATP, 0.3 NaGTP, and 10 phosphocreatine (310 mOsm/L). Neurons were recorded 150-500 μm below the cortical surface. Current clamp recordings were performed with a MultiClamp 700B patch clamp amplifier (Molecular Devices). Current flow out of the amplifier into the patch pipette was considered positive.

### Stimuli

We constructed stimuli consisting of sequences of discrete 20 ms pulses, delivered as either light flashes, auditory clicks or whisker deflections. Each trial was composed of pulses presented at fixed frequencies or following a homogeneous Poisson process with rates ranging from 0.5-20 pulses/s. Fixed frequency stimulation was set at a four second duration per trial while Poisson trials varied in their length. An additional stimulus for model validation was generated as an inhomogeneous Poisson process with a slowly varying rate (time constant of 1 ms) ranging from 0.5-20 pulses/s. The fixed frequency stimulus set was designed to directly measure how the dynamics of the response systematically change with stimulation frequency. The Poisson stimulus sets were used to fit linear/nonlinear models applying maximum likelihood techniques in order to predict the responses to the range of fixed frequencies.

Visual: Full-field monitor flashes were presented monocularly at full contrast on a black screen. All stimuli were generated via the Psychophysics Toolbox (Brainard, 1997; Pelli, 1997) for MATLAB (Mathworks) on a Macintosh (Apple) computer. Stimuli were presented on a calibrated CRT monitor (Sony FDM-520) placed 25 cm in front of the animal’s eyes with a refresh rate of 100 Hz and a spatial resolution of 1,204 × 768 pixels. The mean luminance of the monitor was 40 cd/cm^2^.

Somatosensory: For whisker deflection the principal whisker (trimmed to 10-20 mm) was inserted into a 21G needle attached to a galvanometer servo control motor (6210H; Cambridge Technology Inc., USA) with a matching servo driver and a controller (MicroMax 677xx; Cambridge Technology Inc., USA). A fast-rising voltage command was used to evoke a fast whisker deflection with a constant rise time of 1 ms followed by a 20 ms ramp down signal to prevent an off response for each stimulus. Because of the fixed rise time, amplitude and speed of deflection grow together following a quasi-linear relationship.

Auditory: Click stimuli consisted of 20 ms bursts of broadband noise (5 ms linear rise/fall ramps). They were transduced to analog signals with a high-quality sound card (RME HDSP-9632), attenuated (TDT PA5), and presented to the contralateral ear (TDT EC1). The noise was generated with a spectrum level of −50 dB/sqrt(Hz), and had a bandwidth of 60 kHz. For acoustic calibration, pure tones were used. Typically, a pure tone at 0 dB attenuation produced a sound level of 100 dB SPL, with variations of up to 10 dB across frequency.

### Data analysis

Spikes were identified and removed by passing membrane potential data through a 10 ms median filter or by interpolation (Meir et al. 2018). The mean membrane potential was computed by averaging all trials for each stimulus frequency.

Response amplitude was assessed for each individual pulse (whisker deflection, monitor flashes, or single noise bursts) in the stimulus train. The peak membrane potential was obtained for each stimulus period and the baseline membrane potential at the time of the stimulus pulse was subtracted away to obtain the response amplitude. These responses were then normalized so that the value of the response to the initial pulse was one and a lack of response was considered zero. Adaptation ratios were obtained by dividing the amplitude of the response to the last pulse by the amplitude of the first response (Meir et al. 2018).

As described below, we found that sensory stimulation sometimes evoked a complex bi-or multiphasic response consisting of an initial rapid phase followed by a dip and then a second response phase. We defined cells as containing a multiphasic response by initially identifying the initial peak response that occurs after the stimulus. We then identified the time point following the initial peak at which the response significantly declined from the peak and arrived at a minimal value for 10 ms. We next measured whether a second response depolarization occurs following this minimum by measuring if the response significantly increased after the minimum time point. Those cells and conditions in which response was significantly larger than the trough were marked as multiphasic. Significant differences were determined by a one-sided t-test (P<.05).

Termination responses were defined as significant depolarizations following the cessation of the stimulus. To test for the presence of termination responses we compared the mean membrane potential before the stimulus train to the mean membrane potential 300-800 ms after the final stimulus pulse with a Wilcoxon rank-sum test. These were distinct from the response to the last stimulus as they occurred a few hundred milliseconds after the last stimulus pulse. Termination response amplitudes were defined at the peak of the trial-averaged response. Latencies were defined as the time from the last stimulus pulse to the peak of the termination response.

### Modeling

We fit our model to all Poisson trials (excluding the repeated Poisson noise stimulus). For each trial, we fit the voltage recorded from 50 ms before the stimulus window onset to 1500 ms after the stimulus window offset. Because the adaptive response dynamics we modeled occurred on timescales larger than 10 ms, we downsampled the median-filtered voltage to 10 ms bins.

We modeled the voltage as a sum of linear-nonlinear subunits. Our approach is similar to previous models of spiking activity in the lateral geniculate nucleus (McFarland et al., 2013) and retina (Freeman et al., 2015; Maheswaranathan et al., 2018). The bank of nonlinear subunits could approximate the input received from distinct presynaptic sources (such as excitatory and inhibitory neurons) which are rectified by synaptic transmission. For time step t on trial i, the voltage is modeled as:

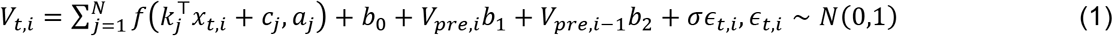

The stimulus before time t is given by the vector *x_t,i_*. The N linear filters are the vectors *k_j_*, and the baseline level for each subunit is *c_j_*,. The nonlinearity is a variation on the logistic function:

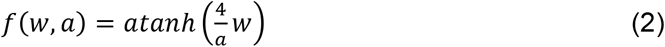

where a is the subunit’s scale, which was restricted to be greater than 1. This formulation gives *f*(0, *a*) = 0 and the derivative *f_w_* (0, *a*) = 1 so that the filters can roughly be viewed in units of mV/pulse, regardless of the scale parameter a.

The model accounts for slow drift in baseline voltage occurring over trials by incorporating the voltage in a 400 ms window occurring 450-50 ms before stimulus onset. The value *V_pre,i_* is the lower 5th percentile value of the voltage in that window as an estimate of the baseline voltage. We also use the baseline estimate from the previous trial, *V*_*pre,i*–1_, to enhance this estimate. The *V_pre_* estimates are weighted by *b*_1_ and *b*_2_. The final baseline term is *b*_0_, which is constant across all trials and times. The noise variance is *σ*^2^.

To reduce the model complexity and promote smoothness in the linear filters, we parameterized the linear filters using a raised cosine basis (Pillow et al., 2005, 2008) of the form:

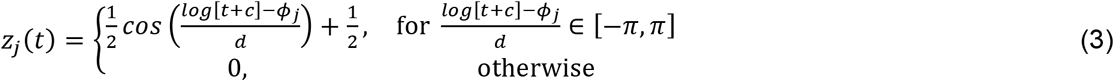

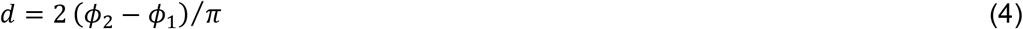

The *ϕ_j_* are spaced linearly from *ϕ*_1_ = log(*t_start_ + c*) to *ϕ_M_* = *log*(*t_end_* + *c*). For V1 and S1 cells, we used M = 16 filters with *t_start_* = 0.01s, *t_end_* = 2*s*, and *c* = 0.3. For the A1 cells, we used M = 20 basis functions with *t_start_* = 0.01s, *t_end_* = 2*s*, and *c* = 0.1 to account for fast timescale responses. For fitting, the basis was orthonormalized.

We placed an independent Gaussian prior on each term in the filter parameters and the history terms:

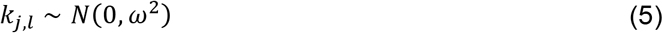

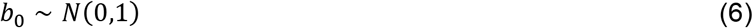

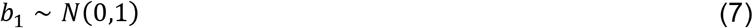

We set *ω*^2^ = 5^2^. This choice of prior regularizes the filter estimates by shrinking the filters towards 0 and keeps the per-pulse deviations in voltage within a biophysically realistic range: a priori, the mean maximum deviation per pulse for a single filter is 4.44 mV with a standard deviation of 1.58 mV for the V1 and S1 bases. Similar results were achieved for different choices of the shrinkage parameter. The other prior used was for the noise term 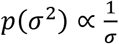. Uniform priors were placed over *b*_0_ and *c_j_*.

We obtained maximum a posteriori (MAP) estimates of the model parameters using a trustregion algorithm. Because this nonlinear model is not convex, we fit the model to each cell 1000 times using random initialization points and selected the fit with the largest log posterior value. The initial conditions were generated according to the distributions:

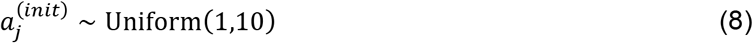

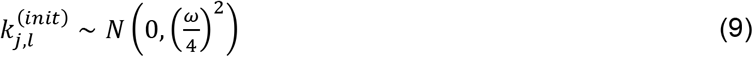

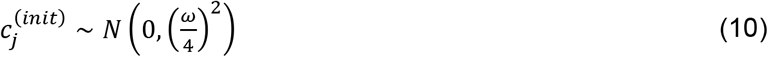

The linear terms *b*_0_, *b*_1_,and *b*_2_ were set to the maximum likelihood estimate computed by normal least squares keeping the other parameters fixed. Similarly, *σ*^2^ was set to the MAP estimate given all other parameters.

We evaluated model performance by predicting the voltage recorded in response to stimuli that had not been used for model fitting. For those stimuli, we tested the model’s ability to predict the average voltage recorded in response to the stimulus instead of predicting single trials. We evaluated model performance using the Pearson’s correlation coefficient between the true and predicted voltages and the percent variance explained:

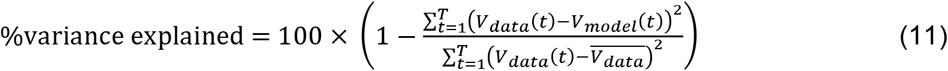

To analyze the model response to frequency changes, we ran model simulations (Fig. 7) composed of stimuli which two stimulus frequencies were presented. The initial frequencies ranged from 1 to 15 Hz and were then changed to a new value between 1 and 15 Hz after 3 seconds. For these simulations, the stimulus pulse times were randomized. Each combination of stimulus frequencies was simulated 5000 times using distinct pulse times and these resulting model outputs were averaged. Frequency transition responses were calculated as the mean 200 ms around the peak of the response. The mean steady state amplitude 200 ms before the transition was then subtracted from this value to obtain the transition response amplitude.

## Results

To examine the adaptive properties of sensory neurons across modalities, we constructed a stimulus set consisting of temporal sequences of constant amplitude, discrete punctate pulses: monitor flashes, auditory clicks or transient whisker deflections. These temporal sequences were either Poisson point processes or fixed-interval trains presented at multiple frequencies. We recorded the membrane potential responses in three regions of mouse sensory cortex (A1: n = 9, S1: n = 14; V1: n = 11) using whole-cell recordings.

In response to temporal sequences composed of fixed intervals, the membrane potential exhibited a number of dynamical properties that appear to be generally conserved across sensory cortex (Fig 1). First, increasing stimulus frequency entailed more adaptation, leading to a systematic reduction in response amplitude towards the end of the train. Second, we observed that the reduction in stimulus-evoked response when stimulus frequency increases is accompanied by a systematic shift in the complexity of the response to individual pulses: at low frequencies individual responses are long-lasting and multiphasic, whereas they become briefer and monophasic at higher frequencies. Finally, we found that a response occurs at the cessation of a high frequency stimulus, which we term a termination response. Neurons across modalities varied in the degree to which they expressed these properties to fixed interval stimulation, as detailed below, but these motifs persisted across our sample database.

**Figure 1.**
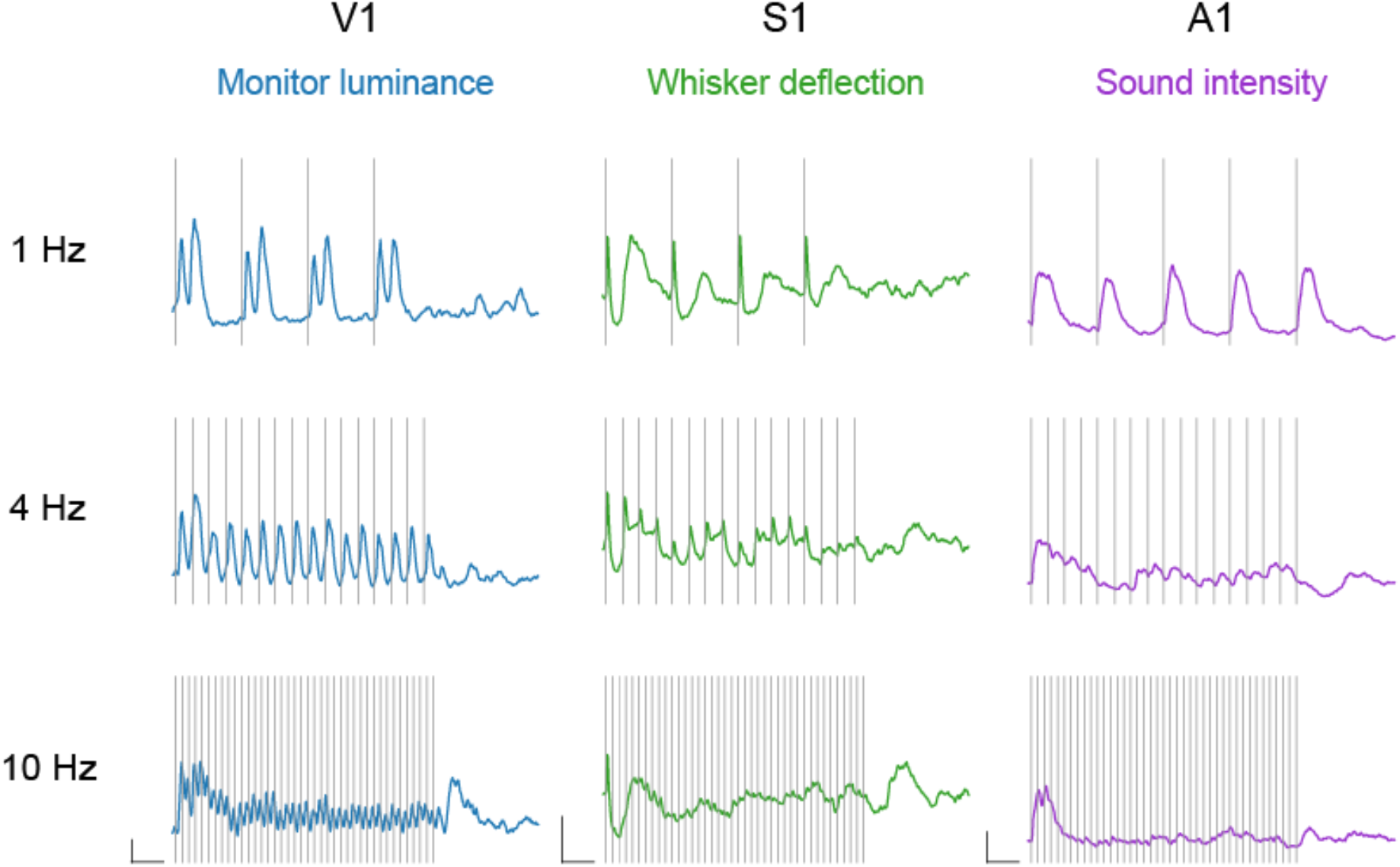
Fixed-interval stimulus responses across sensory cortex. Example V1 (blue), S1 (purple) and A1 (green) membrane potential responses to fixed-interval stimuli delivered at 1 Hz, 4 Hz and 10 Hz. Scale bars indicate 5 mV change in membrane potential on the y-axis and 500 ms duration on the x-axis. Gray bars represent an individual pulse in the stimulus train.

One way these features could arise is if the state of the system changes as a result of being driven by stimuli with different statistics (Garrido et al. 2009). For instance, an increase in stimulus frequency could alter response time constants, resulting in a more monophasic response to the individual stimuli. Such a possibility could arise by adaptation recruiting slow inhibitory inputs (Dealy and Tolhurst, 1974). This would render any attempt to predict responses across stimulus frequencies from a simple fixed model fruitless, as model parameters would need to be adjusted with the stimulus statistics. Alternatively, varying stimulus statistics may evoke different components of a fixed but complex response. In this case, changes in response due to altered stimulus statistics could be modeled as an emergent property of the combination of a single set of sensory filters.

To determine whether a single set of filters can account for the responses seen across stimulus frequencies, for the same cells we additionally recorded membrane potential responses to Poisson pulse trains that varied widely in their rates, where the amplitude and shape of each stimulus was identical to those used for the fixed-frequency stimulation. We then fit these data within a linear-nonlinear modeling framework. Our model was composed of a bank of linear filters that were followed by output nonlinearities (see Methods, Fig. 2B). The outputs of the linear-nonlinear subunits were summed to give the estimated membrane potential. This architecture provides flexibility in allowing multiple contributions to membrane voltage, or “subunits”, with independent nonlinearities. We used maximum *a posteriori* methods to estimate the parameters of this linear-nonlinear sub-unit model (Eq. 1) using each cell’s responses to Poisson trains with a wide range of mean stimulation rates. Because the pulse stimuli are non-Gaussian, we used maximum likelihood fitting instead of an approach like spike-triggered covariance (STC) which is most appropriate for Gaussian stimuli (Brenner et al. 2000; de Ruyter van Steveninck & Bialek, 1988; Schwartz et al., 2006; Park et al., 2013; Aljadeff et al., 2017).

**Figure 2.**
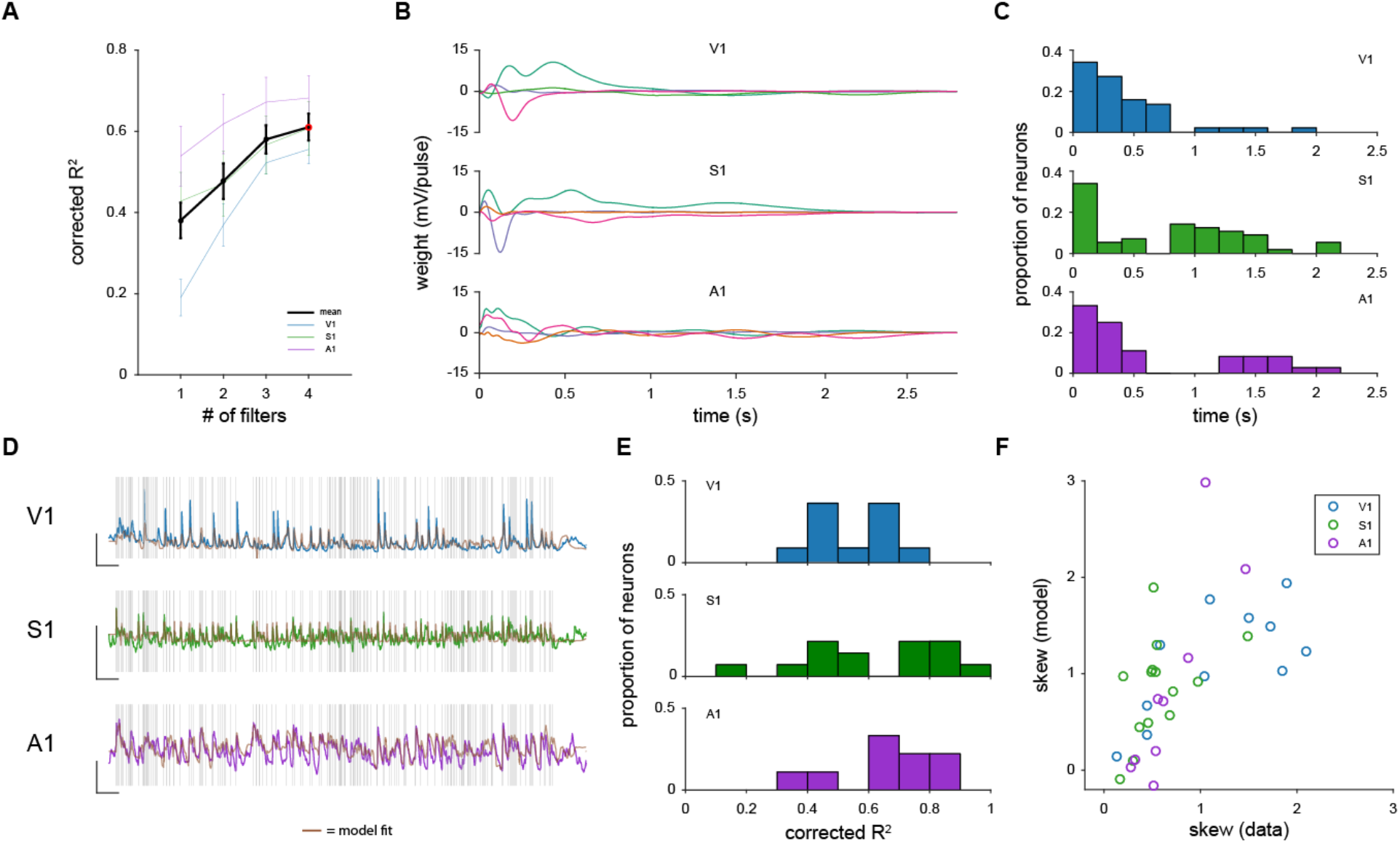
Modeling framework. (A) Corrected R2 values to a withheld stimulus generated by an inhomogeneous Poisson process by number of filters included in the model. Data are mean +/− standard error (B) Example filters for individual neurons for all three sensory systems. (C) Histograms of filter timescales across the population. (D) Model fits obtained from the example filters shown in (B) to a fixed Poisson-process stimulus. Scale bars are 10 mV on the y-axis and 1 second on the x axis. Brown traces represent model fits. Gray bars represent individual stimulus pulses. (E) Histogram of variance accounted for to the fixed Poisson stimulus for V1, S1 and A1. (F) Scatterplot comparing the skew of the responses to the fixed Poisson-process stimuli for the data and the model simulations.

We first examined how the number of subunit filters included affected the model’s ability to capture response dynamics to Poisson stimuli for individual neurons. We assayed the model performance by examining how well it could account for the average response to a separate (held-out) Poisson sequence that had been repeated many times (Fig 2D.) Across V1, S1 and A1, models composed of 4 filters accounted for a significant amount of the explainable response variance to the repeated Poisson sequence (R^2^: V1 = .47 +/− .04; S1 = .4 +/− .06; A1 = .50 +/−..07; values indicate mean +/− SE across neurons), demonstrating that this modeling framework can largely account for sensory responses in cortex. To estimate the fraction of explainable variance the model accounts for, we employed a method developed by Sahani and Linden (2003) which takes into account the number of stimulus repeats and the variation between trials (see Sahani & Linden, 2003; Mohanty et al. 2012). This correction factor only modestly increased the variance accounted for by our model (Fig 2E; corrected R^2^: V1 = .56 +/− .04; S1 = .61 +/− .07; A1 = .68 +/− .06). Increasing the number of filters systematically improved the model’s performance, although performance increases from using more than 4 filters were minimal (Fig. 2A). Beyond overall performance, a model composed of a single filter failed to account for specific features of the adaptive response to fixed frequency stimuli, as detailed in the following sections. To standardize our analyses across neurons and modality we fixed the number of filters used to 4.

To characterize how the model is able to match the responses to Poisson sequences we examined the properties of the subunits used to generate the model. We extracted the temporal envelope of each subunit filter by performing an autocorrelation and quantifying the time over which it was above 0.2 (Shelhamer, 2007). For individual neurons these subunit timescales could vary from 60 ms to over 2 seconds. Most subunit filters had short envelopes, forming an exponential distribution of timescales (Fig. 2C). Each neuron had filters that varied in time scale. The median difference between the fastest and slowest time scale was over 1 second. The average ratio of the time of the longest and shortest filter for each cell was 8.48 +/− 1.24 (geometric mean). Because of these subunit time scales we set a maximum filter length of 2.5 seconds for each subunit of our model.

We next sought to determine how well the model could account for a membrane potential skew. We chose skew because this metric of the response distribution encapsulates the degree of neuronal selectivity, particularly when a broad stimulus range is employed. Those neurons that respond to specific stimulus conditions have higher skews, whereas those that respond more broadly have lower skews (Ringach and Malone, 2007). The model was largely able to predict the skew of the membrane potential response to the left-out Poisson sequence across sensory areas (Fig 2F; Correlation coefficient of skew (r): V1 = .70; S1 = .46; A1 = .84)

We tested whether the subunit model, fitted from noise sequences, was able to capture the membrane potential fluctuations observed in response to fixed interval sequences, (Fig. 1A). The subunit model provided predictions that were highly correlated with the actual responses of the neurons (corrected R^2^: V1 = .68 +/− .04, S1 = .73 +/− .05, A1 = .83 +/− .05). These high correlations demonstrate that in general the model predicts the membrane potential responses observed in the fixed interval data.

We next sought to focus our analysis on specific components of the adaptive response. We examined whether the model could recapitulate the three prominent adaptive aspects of the responses highlighted above: 1) the decay of response amplitude when increasing stimulus frequency, 2) the shift in complexity of a single-pulse response from biphasic to monophasic as stimulus frequency increases, and 3) the presence of a termination response following the termination of a high frequency stimulus train (Fig. 1).

### Response attenuation with stimulus frequency

A prominent feature of sensory adaptation is that the degree of response attenuation is linked to the rate of sensory stimulation: higher frequency stimulation leads to stronger amplitude attenuation (Chung et al., 2002; Khatri et al., 2004; Kheradpezhouh., 2017; Martin-Cortecero & Nuñez, 2014). We find this common pattern in our V1, S1 and A1 responses (Fig. 3A). The responses of neurons at 1 Hz stimulation were only weakly attenuated, whereas the responses to 10 Hz stimuli were strongly attenuated. To determine the degree of attenuation to each fixed-frequency stimulus, we quantified the membrane potential response to each pulse in the train (Fig 3B, see methods). The peak membrane potential response to each pulse in the stimulus train was obtained and normalized to the value at the time of the stimulus pulse. In these example neurons we find that response amplitude systematically declines as a function of stimulus frequency and the location of the pulse in the train. These example neurons reflect the typical responses found in each modality, in which low frequency stimulation evokes little response attenuation and high frequency stimulation evokes large attenuation.

**Figure 3.**
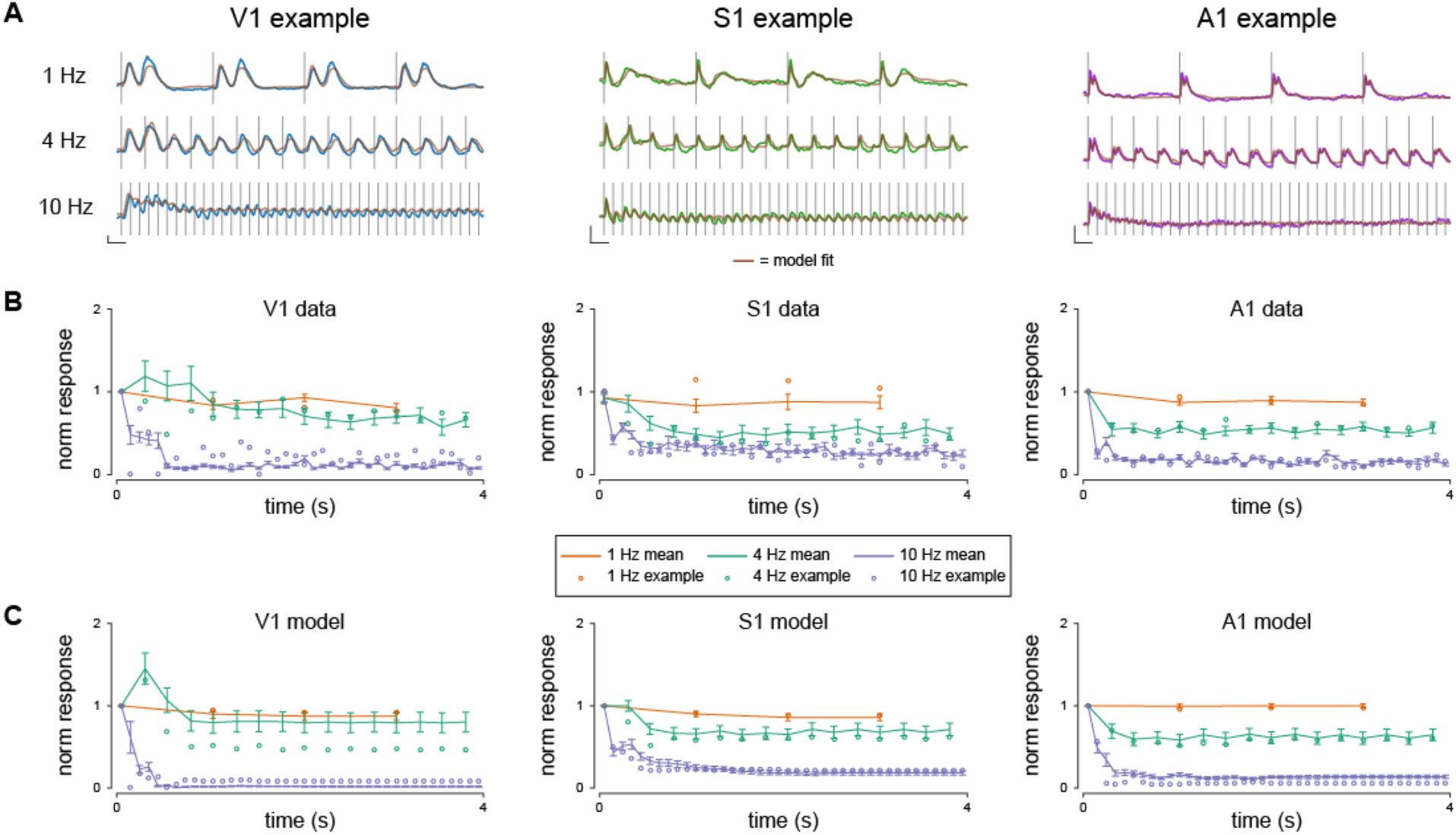
Response attenuation increases with stimulus frequency. (A) Example V1, S1 and A1 membrane potential responses to fixed interval stimuli delivered at 1 Hz, 4 Hz and 10 Hz. Brown traces represent model fits to these responses. Scale bars indicate 5 mV change in membrane potential in the y-axis and 200 ms duration in the x-axis. (B) Normalized mean peak membrane potential responses to each pulse in a stimulus train at the three frequencies shown in (A). Open circles are data from the example cells in (A). Error bars represent SEM. (C) Same as (B) but for model fits.

To quantify the frequency dependence of this attenuation in our dataset, we computed the adaptation ratio, defined as the ratio of the response amplitude of the last stimulus in our train and the response to the first stimulus, for each stimulus frequency (Meir et al. 2018). Although there was considerable variability in adaptation across and within sensory systems, a pattern emerges when looking at the population. For 1 Hz, the adaptation ratio was similar across V1, S1 and A1 neurons (Fig. 4). As frequency increased, these adaptation ratios systematically declined, such that at 10 Hz, the adaptation ratios were much closer to 0. Neurons attenuate in a similar manner within a modality, and the same trend of greater attenuation with higher stimulation frequency is observed across modalities. Hence, under comparable experimental conditions, response attenuation follows a similar pattern for these 3 sensory modalities.

We tested whether our model could account for these findings by simulating responses to the 1 Hz, 4 Hz and 10 Hz stimuli for each cell in our dataset (Fig. 3A, brown traces) using the filters that were obtained from Poisson stimulation. We then performed the same analysis on these model fits to determine whether they exhibit the same response attenuation (Fig. 3C). Adaptation ratios of model fits were highly correlated with their data counterparts (r = .80 (V1), .68 (S1), .95 (A1), see figure 4). These data indicate that our model is able to capture the stimulus frequency dependence of response attenuation across three areas of sensory cortex. Note that not only is the model able to capture the broad response attenuation observed in our dataset, but it predicts differences in response attenuation across modalities. For example, we find that S1 neurons attenuate less at 10 Hz than either A1 or V1 neurons, which matches model predictions.

**Figure 4.**
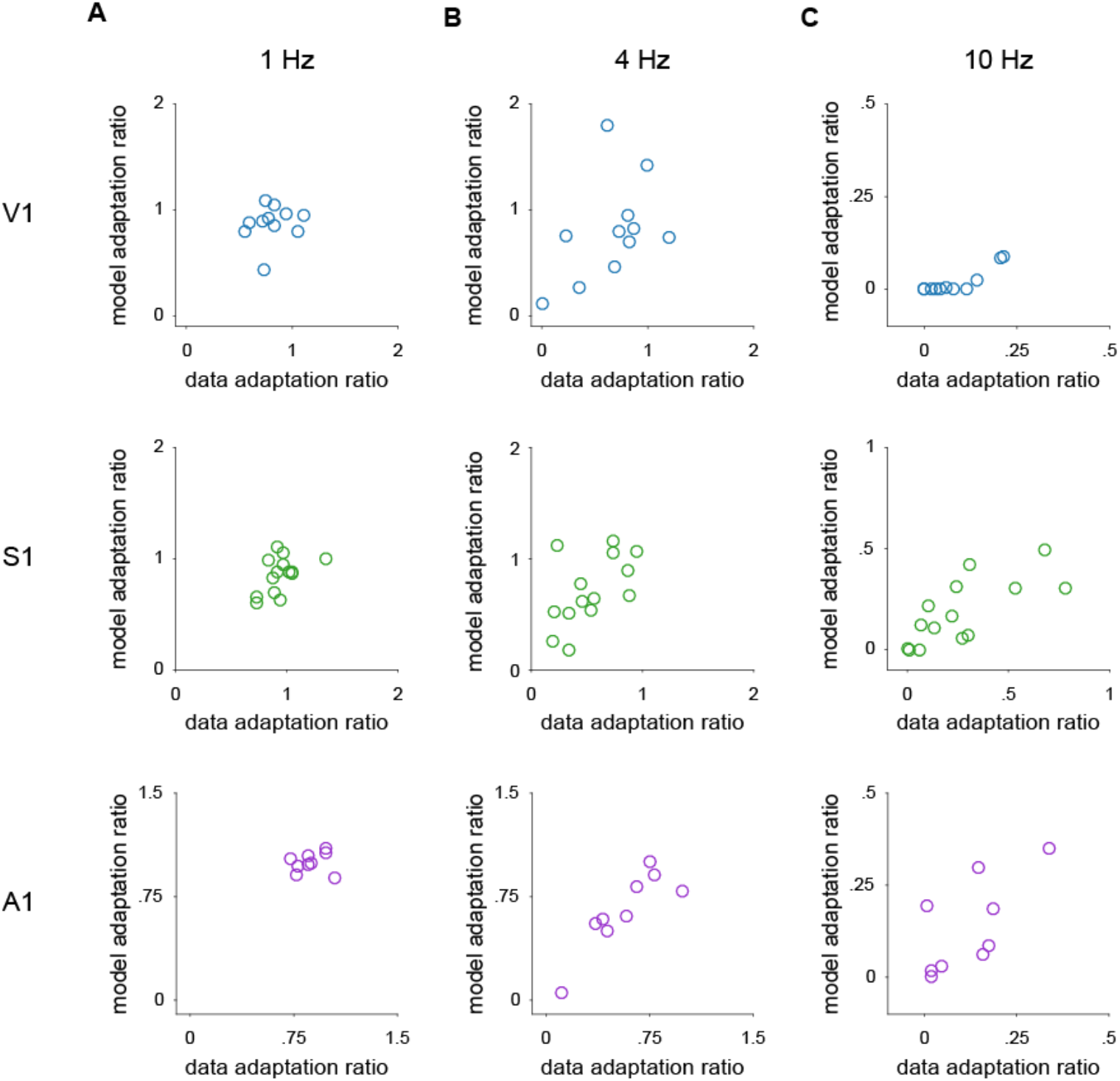
Model fits account for response attenuation adaptation ratios. (A) Scatterplot comparing the adaptation ratios of neurons for the data (x-axis) and model fits (y-axis) in response to a 1 Hz stimulus for V1, S1 and A1. (B) Same as (A) but for 4 Hz stimulus. (C) Same as (A) but for a 10 Hz stimulus.

### Change in response complexity with stimulus frequency

The second adaptive feature we noticed was that the shape of the membrane potential response is often altered by stimulus frequency. Visual inspection revealed a characteristic shift in the complexity of the response to individual stimuli as stimulus frequency increased in a large proportion of visual and somatosensory neurons (Fig. 5A). At lower frequencies (< 4 Hz), complex biphasic responses were common in S1 and V1, as has been previously reported in rodents and humans (Funayama et al., 2016; Funayama et al. 2015; Sachidhanandam et al., 2013). We found, however that these multiphasic responses shifted to simple, monophasic responses as stimulus frequency increased to 4 Hz. We classified whether the responses were significantly multiphasic by determining whether responses followed a pattern of initial depolarization, a decline in membrane potential followed by a second depolarization that is significantly larger than the decline. The multiphasic responses apparent at low stimulus frequencies follow this pattern, responding with a significant depolarization following the dip. Low-frequency stimuli evoked membrane potential responses that were deemed multiphasic for the majority of V1 (73%) and S1 (93%) neurons (Fig 5B), but only the minority of A1 neurons (44%). A1 neurons exhibited a somewhat different type of biphasic response, with much shorter latencies between the two components of the response (Figure 5A, bottom trace). When the stimulus rate was increased to 4 Hz, the responses of V1, S1 and A1 neurons became less multiphasic and none exhibited multiphasic responses.

**Figure 5.**
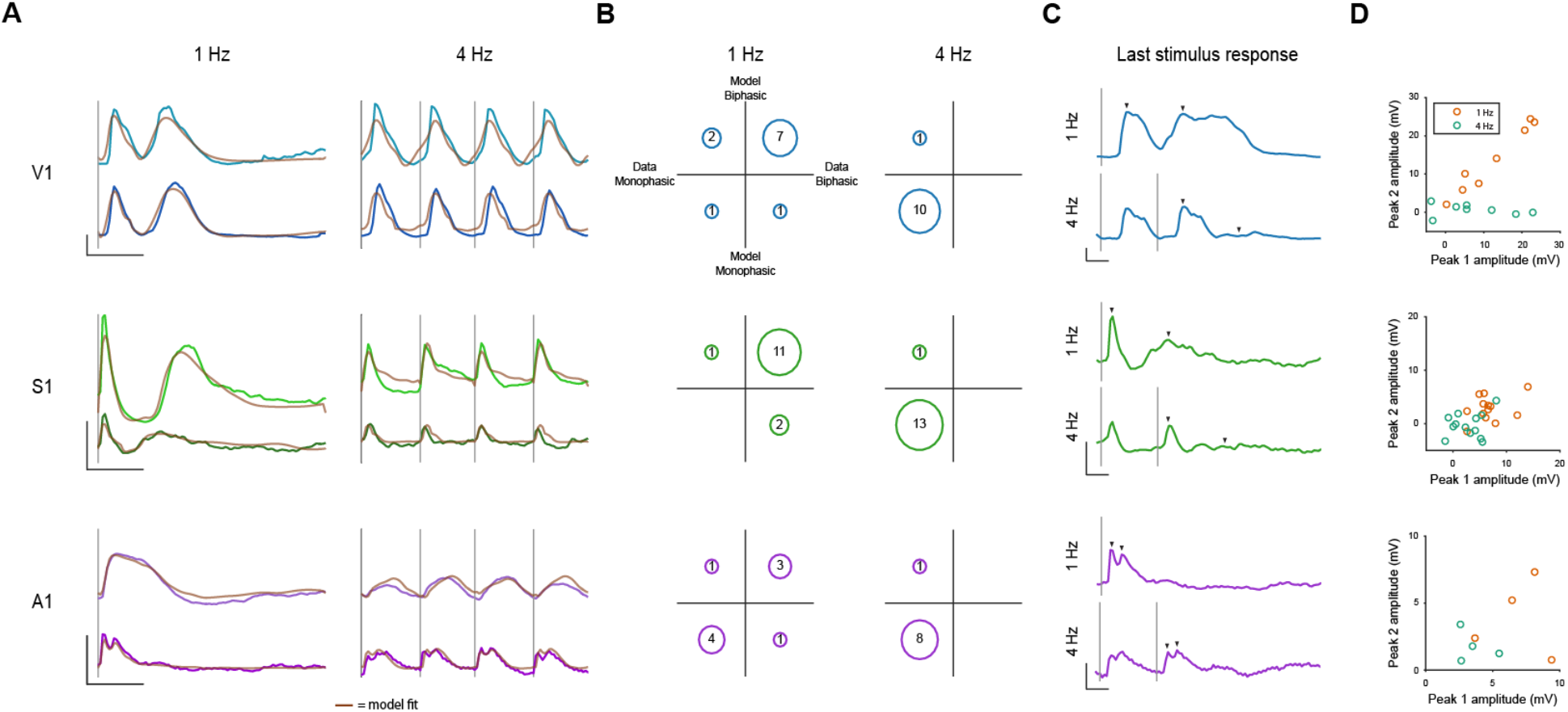
Change in response complexity with increasing stimulus frequency. (A) Example V1, S1 and A1 membrane potential responses to fixed interval stimuli delivered at 1 Hz, 4 Hz. Two example cells per sensory system are shown. Brown traces represent model fits to these responses. Scale bars indicate 10 mV change in membrane potential in the y-axis and 200 ms duration in the x-axis. (B) Count of neurons and model fits whose responses are significantly biphasic in V1, A1 and S1. The x-axis categories data as monophasic (left) or biphasic (right). The y-axis categorizes model fits as monophasic (bottom) or biphasic (top). (C) Example membrane potential responses to fixed interval stimuli delivered at 1 Hz, 4 Hz. Traces show the end of the stimulus train. Arrows indicate identified peaks. Scale bars indicate 5 mV change in membrane potential in the y-axis and 100 ms duration in the x-axis. (D) Scatterplots of peak amplitude for both peaks of the biphasic response. Peak locations were determined from the 1 Hz trace and were then measured on the 4 Hz stimulus from the time of last stimulation.

Not only did the response complexity change with stimulus frequency, response duration also shifted with stimulus frequency. We quantified response duration by measuring the envelope of time over which responses deviated from the baseline membrane potential (Methods). We examined the response stimuli that started 2 seconds after the stimulus train began to be sure that the cell had adapted to the stimulus frequency. Responses to 4 Hz stimuli were considerably shortened in V1 and S1 neurons (V1 = 126 ms +/− 39, S1 = 101 +/− 65 ms) relative to response to 1 Hz stimuli (V1 = 505 +/− 138 ms, S1 = 435 +/− 237). Responses for A1 neurons, however, were similar between the two frequencies (1 Hz = 158 +/− 101 ms, 4 Hz = 158 +/− 56 ms).

We next sought to determine whether the subunit model could account for these response dynamics (Fig 5A; brown traces). As with the recordings of the membrane potential, we measured whether model responses were significantly multiphasic. The model successfully recapitulated the multiphasic responses in V1, S1 and A1 at low frequencies (V1 = 82%, S1 = 86%, A1=44%) as well as the shift to monophasic responses at higher frequencies (V1 = 9%, S1 = 7%, A1=11%; Fig 5B). Furthermore, the results revealed that, in general, the neural responses and their model fits received the same classification (Fig 5B). The model also captured the large shift in response duration in V1 (1 Hz = 507 +/− 118 ms, 4 Hz = 146 +/− 27) and S1 (1 Hz = 507 +/− 227, 4 Hz = 151 +/− 58 ms) and the weak shift in A1 responses (1 Hz = 216 +/− 132 ms, 4 Hz = 137 +/− 58 ms). Across modalities, the subunit model resulted in response durations that were correlated to those measured in V1, S1 and A1 (r: V1 = .92; S1 = .77; A1 = .70).

One possibility is that responses seemed monophasic at higher frequencies because the biphasic portion of the response was interrupted by the next stimulus pulse. To control for this, we focused our analysis on the last pulse in the stimulus train (Fig 5C). We first measured the mean amplitude 10 ms around the first and second peak of biphasic responses using the indices we obtained previously. The amplitude of the two peaks was highly related in V1 neurons (Fig 5D, r = .98) where the amplitude of the two peaks was generally on the same scale (mean amplitude: first peak 12.3 +/− 3.2 mV; second peak 13.6 +/− 3.1; ttest: p > .05). S1 responses tended to have weaker second peaks (first peak = 6.8 +/− 0.9 mV, second peak = 2.8 +/− 0.6 mV; ttest, p < .001) which resulted in a weaker correlation (r = .36).

We then used these indices to measure the amplitude evoked by a 4 Hz stimulus at the end of the stimulus train. As previously shown (see previous section) the amplitudes of the first peak were smaller in the 4 Hz condition as compared to the 1 Hz condition (V1 = 7.5 +/− 3.4 mV; S1 = 2.9 +/− 0.8 mV; ttest, V1: p = .02; S1: p=.003). Using the latencies obtained from the 1 Hz responses, we measured the amplitude of the “second peak” of the 4 Hz responses to the last stimulus in the train. As expected, amplitudes were near zero, indicating a lack of a biphasic response (V1 = 0.6 +/− 0.5 mV; S1 = −0.3 +/− 0.6 mV). Furtermore, the amplitude of the first 4 Hz peak exhibited weak and non-significant correlations with that of the second (r: V1 = −.25, p>.05; S1 = .23, p>.05). These results indicate that 4 Hz responses truly shift to a monophasic shape and are not an artifact of our protocol and analysis. A1 data are not summarized here due to the low number (4) of biphasic neurons, but data are shown in Fig 5C,D (bottom row).

### Termination responses

Paradigms employing periodic stimuli have consistently found a large response at the end of a stimulus train or when a stimulus is omitted or changed (Bullock et al., 1990; Hamm & Yuste, 2016, Karamursel & Bullock, 1994; Li et al 2017; Näätänen et al., 1978; Schwartz et at. 2007). These phenomena have been reported at multiple levels of analysis including EEG (see Näätänen et al., 1978) and in single unit recordings (see Schwartz et at. 2007) in multiple sensory systems. Although the terms (mismatch negativity, omitted stimulus response, echo response, etc.) and the underlying mechanisms of these responses may differ, their descriptions and interpretations share a number of common features (Bullock, Karamursel & Hofmann, 1993; Schwartz et al., 2007; Stefanics, Kremlacek & Czigler 2014).

We also found transient membrane potential depolarizations following the termination of the high frequency stimulus trains across the three sensory modalities and term these deflections “termination responses” (Figure 6A). These termination responses are easily distinguished from a response to the last stimulus of the train as they start hundreds of milliseconds after the termination of the stimulus train. To determine whether these depolarizations were statistically significant, we compared the mean membrane potential 300-700 ms after the response to the last pulse in the stimulus train to the membrane potential prior to sensory stimulation. We found that termination responses were present in the majority of V1 (9/11) and S1 (10/14) cells at a stimulation frequency of 10 Hz. Termination responses were found in a smaller proportion of A1 cells (3/9). While termination responses were common at high frequencies, they were less prevalent at lower frequencies such as 4 Hz (V1, 3/11, S1, 2/14, A1, 0/9, Fig. 1). We quantified the latency and amplitude of the termination responses for all of the intracellular records. The latency of the termination response (defined from the last stimulus to the peak) was generally long (V1 = 367 +/− 79 ms, S1 = 446 +/− 102 ms, A1 = 642 +/− 284 ms, see Methods), and its amplitude was large, on generally on the same scale as the response to the first stimulus (V1 = 17.9 +/− 11.2 mV, S1 = 7.6 +/− 3.5 mV, A1 = 4.4 +/− 3.7 mV, Fig 6B). To determine the role of synaptic input to these termination responses, we performed voltage clamp recording while holding the neuron at the reversal potential of inhibition. Voltage clamp recordings indicate a large excitatory current after the end of stimulation that indicates a synaptic origin for these responses (Fig 6C).

The subunit model successfully accounted for the depolarizations at the termination of high-frequency stimulus trains for those neurons with significant termination responses. Despite predicting the presence of a termination response to the high frequency data, our models consistently underestimated the response amplitude at the end of the stimulus train (Fig. 6A, r = .59, slope = .14). The model did predict the long latency of these termination responses though the individual diversity in termination latency was only weakly correlated to our estimates of termination response latency from our measurements (Fig. 6B, r = .414, slope = .41). Some of the difference in latencies between the model and measurements reflects the difficulty assigning a single latency value to responses that extend and slowly depolarize over hundreds of milliseconds.

We focused our comparison of the termination response on two frequencies that either lacked a termination response (4 Hz) or consistently exhibited a termination response (10 Hz). To expand our analysis to include multiple frequencies that exhibited a termination response, we performed additional V1 recordings with a wider range of higher stimulus frequencies (Fig 6D.) We found that the termination responses exhibit changes in amplitude and latency that are related to the stimulus frequency. In particular, the amplitude of the termination response monotonically increased with stimulation frequency (fig 6E). Not only was there a smooth dependence of termination amplitude on stimulation frequency, we also uncovered a systematic change in termination latency on stimulation frequency such that higher frequency stimuli, to which neurons exhibited the strongest adaptation, resulted in shorter latencies of larger amplitudes (Schwartz et at. 2007). These observations indicate that adaptive processes carry information about changes in the temporal statistics of temporal sequences, that is the termination of the stimulus.

**Figure 6.**
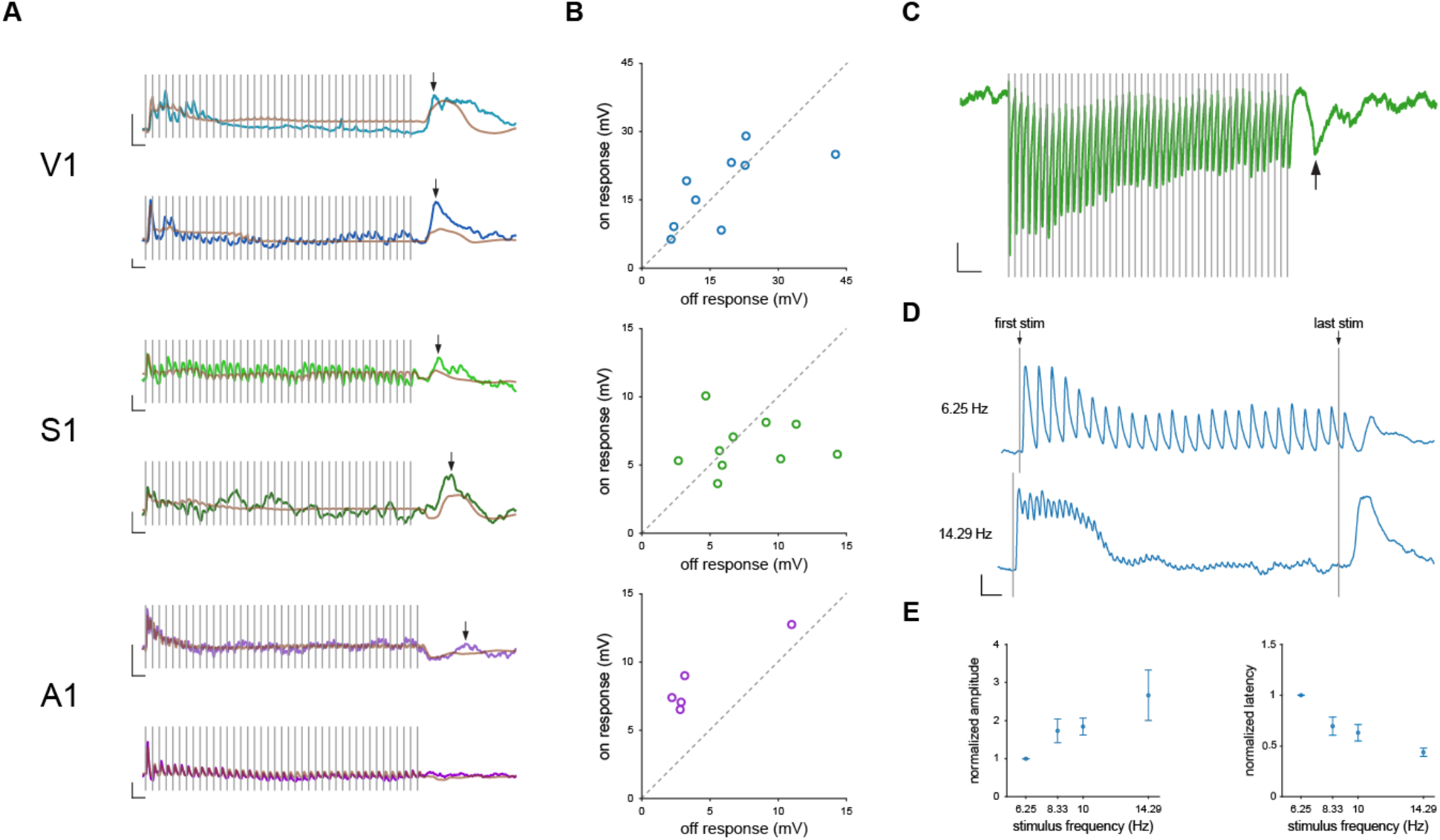
Termination responses. (A) Example V1, S1 and A1 membrane potential responses to fixed interval stimuli delivered at 10 Hz. Two example cells per sensory system are shown. Brown traces represent model fits to these responses. Scale bars indicate 5 mV change in membrane potential in the y-axis and 200 ms duration in the x-axis. Arrow indicates the location of the termination response. (B) Scatter plots comparing the amplitude of the termination response and the on response for V1, S1 and A1. (C) Example S1 voltage clamp recording while the neuron was held at the reversal potential of inhibition. Arrow indicated location of termination response. Gray bars represent individual stimulus pulses. Scale bars indicate a 20 pA change on the y-axis and a 250 ms change on the x-axis. (D) Example neuron membrane responses to two different stimulus frequencies. Scale bars indicate 5 mV change in membrane potential and 250 ms duration. (E) Mean normalized termination response amplitude and latency at four stimulus frequencies. Responses are normalized to the termination response at 6.25 Hz for each cell. Error bars represent the standard error of the mean.

### Signals for changes in temporal statistics

To explore how adaptation might shape signals related to a change in the temporal statistics of our stimuli we explored two distinct paradigms in which the sequences of punctate stimuli are varied. Because the model predicted responses to the absence of stimuli at the end of a sequence, we reasoned that the model may also predict responses when stimuli are removed during a stimulus sequence. Indeed, we find that our models consistently predict a response to an omitted stimulus, or an enhanced response to subsequent stimuli (Fig 7A). This model prediction is exhibited in the responses of individual neurons when presented with the same stimulus train (Fig. 7B). Indeed, such omitted stimuli are particularly salient perceptually (Näätänen, 2018), mirroring the changes in activity we observe and predict from the model.

We next explored how model neurons respond to changes in the frequency of stimulation *f*, reasoning that signals related to the degree of frequency change should be present. We simulated responses from model neurons using filters fit from data to stimuli composed of an initial frequency (F1) which ranged between 1 and 15 Hz. Once the neuron had adapted to the initial frequency, the frequency switched abruptly to a new value which also ranged between 1 and 15 Hz. (F2, see methods). Frequency increases resulted in changes in membrane potential depolarization with amplitudes that were related to the size of the frequency increase (Fig 7C). We noticed two distinct response types in our dataset. When holding F1 constant and varying F2, some cells exhibited a transient response depending on F2 which then quickly settled to a common steady state level (Fig 7C, neuron 1). Other neurons exhibited a similar depolarization, but rather than repolarizing, maintained an elevated membrane potential for the duration of the stimulation, continuously conveying information about the value of F2 (Fig 7C, neuron 2). When holding F2 constant while varying F1, similar patterns emerged (Fig 7D). Surprisingly, in some neurons, although they settled to a similar response amplitude independent of F1, they responded with distinct amplitudes when the frequency transitioned to the common F2 (Fig 7D, neuron 3), showing that information about F1 was retained in the system’s state even though it was not manifest in the membrane potential. Furthermore, the effects of the distinct F1 rates lingered for nearly 2 seconds while responding to the common F2 (Fig 7D, neuron 4). These transition responses suggest that stimulus history can affect dynamics at the membrane potential level for a long time, on the order of seconds.

How might stimulus history and the amplitude of the transition response be related? One possible model is that the change in overall frequency (F2-F1) determines the size of the response, which we term the linear model. In this case, the responses scale linearly with ΔF, regardless of the initial frequency. Another possibility is that the transition amplitude follows Weber’s law. In the Weber model, the response to frequency change scales with respect to value of F2 divided by F1 (F2/F1). In this case, the response to a frequency change depends on Δlog(F), regardless of the initial frequency (logarithmic model).

To determine whether our data follow the linear or logarithmic model, we quantified the transition response amplitude to each frequency change (see methods). We focused our analysis on values of F1 that ranged from 5-10 Hz and F2s that increased F1’s value by 1-5 Hz. We then performed a least square fit to each model. We obtained the slope (mln) and y-intercept (bln) of the best fit line to the transition response amplitudes from an initial frequency of 5 Hz on a linear (F2-F1) x-axis. The linear model was defined using the following equation (Fig. 7E):

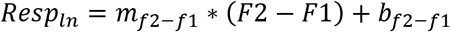

A similar procedure was then performed to obtain the Weber’s law slope (mwb) and y-intercept (bwb) using the same response amplitudes as before on a Weber-like (F2/F1) x-axis. The logarithmic model was defined using the following equation (Fig. 7F):

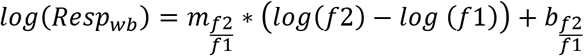

These two examples show that ‘Neuron 2’ exhibited linear responses, whereas ‘Neuron 1’ showed more logarithmic-like responses. To classify each neuron as following either the linear model, logarithmic model or neither, we used a correlation analysis which removes shared correlation (Fig. 7F. Movshon et al., 1986). We calculated partial correlations using the actual and predicted responses from each model to each stimulus condition using the following equations:

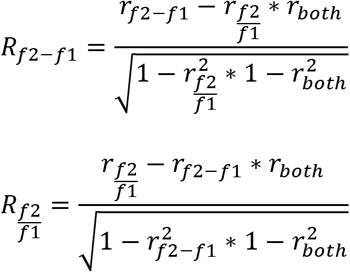

where r_f2-f1_ and r_f2/f1_ are the correlation between the respective model predictions and responses and r_both_ is the correlation between the two model predictions. We then calculated Fisher z-transformed partial correlations using the following equations:

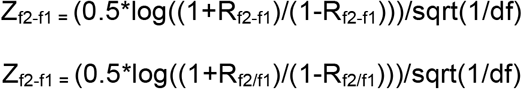

This analysis reveals that a sizeable proportion of the fitted neurons follow either the logarithmic or linear model (29/34: V1: 10/11, S1: 11/14, A1: 8/9). Of these, 59% (V1: 80%, S1: 36%, A1: 63%) significantly followed the logarithmic model. Note however, that we found neurons in each sensory modality that carry signals related to linear changes in frequency as well as relative changes in frequency and that these distinctions reflect differences in our population in degree rather than category.

**Figure 7.**
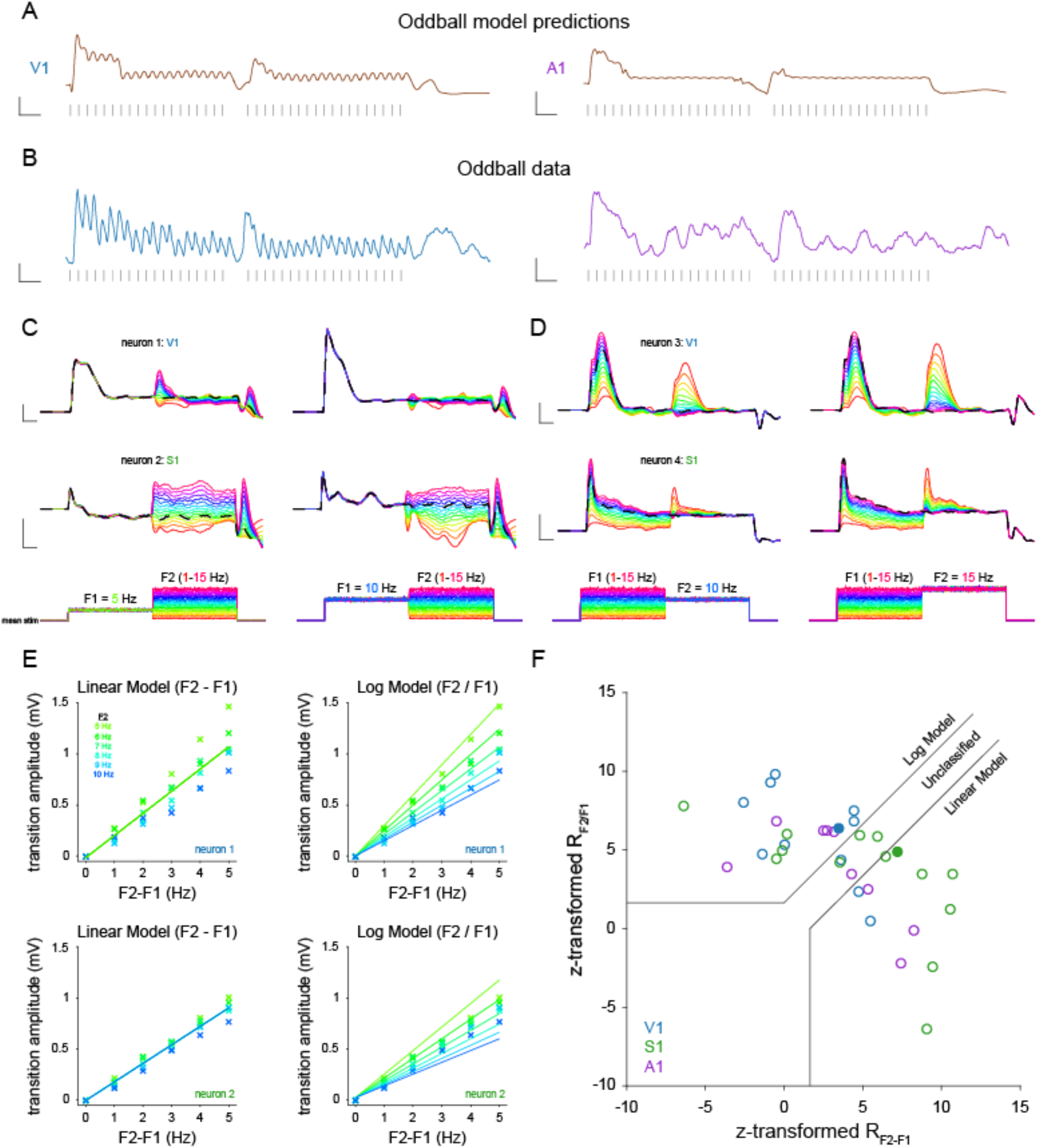
Responses to frequency transitions. (A) Model predictions for two neurons (V1, A1) for an oddball stimulus. One or two stimulus pulses was removed in the middle of the stimulus train. Scale bars indicate a 5 mV change on the y-axis and 250 ms duration on the x-axis. (B) Membrane potential data for the oddball stimulus. Scale bars indicate a 5 mV change on the y-axis and a 250 ms duration of the x-axis (C) Model simulations for two cells. Initial mean stimulus frequency was 5 Hz (top trace) or 10 Hz (bottom trace) for all traces. This then transitioned to all frequencies between 1 and 15 Hz. Dotted line indicates no frequency transition. Scale bars indicate a 2 mV change on the y-axis and a 500 ms duration on the x-axis. (D) Same as (C) but the second frequency was held constant and the first frequency was changed (E) Model predictions for a linear and weber frequency transition signal. Solid bars indicate prediction, x’s indicate data. (F) Scatter plot of z-scored partial-correlations. Solid lines indicate significance at 95%. Filled circles are example cells from (C).

## Discussion

We examined adaptation in visual, somatosensory and auditory cortex using a common stimulus framework to uncover their adaptive response properties. We used random Poisson input trains to develop models for responses as a sum of independent linear-nonlinear subunits, and found these models to be surprisingly effective in predicting the adaptive effects seen in our responses. Our results extend upon previous studies that have shown that many nonlinear properties of neural responses can be captured by summing independent subunits (Ozuysal & Baccus, 2012; McFarland et al., 2013; Freeman et al., 2015; Vintch et al., 2015, Harper et al., 2016). We find that for each neuron, a fixed set of four filters spanning multiple timescales accounts for response dynamics across stimulation frequency, demonstrating that a single static model accounts for responses under distinct sensory contexts. Therefore, these adaptive changes need not reflect a change in the state of the system but rather can accounted for by modeling adaptation as an emergent property of fixed, but complex, neural responses.

We focused on three components of the adaptive sensory response: decreased response amplitude with increasing stimulus frequency, a shift in response complexity as stimulus frequency increases, and a termination response at the cessation of a high frequency stimulus. We employed common stimulus sets and analysis tools to study the adaptive properties of these response features across modalities, allowing us to demonstrate which are common across modalities and which are more specific. We find, for example, that decreases in response amplitude with increased stimulus frequency are shared across modalities but vary in degree. The frequency dependence of response complexity was common to both S1 and V1. While individual A1 neurons had distinct response dynamics, we did not find a frequencydependent shift in those dynamics. Finally, all modalities demonstrated a termination response following the end of stimulation in at least some of the neurons.

All three of these response characteristics emerge also from our model. We thus demonstrate that a general model built from responses to stochastic stimuli, with a broad statistical range, can dissect complex neural responses to fixed-interval stimuli into simpler components to reveal general features of sensory adaptation. The emergence of these adaptive features relies on a model composed of multiple filters. When a model composed of a single filter is used, overall model performance sharply declines. Furthermore, the specific adaptive responses to fixed frequency seen in our data are absent from single filter model responses. The use of multiple filters with associated nonlinear transformations accounts for the ability of our model to generate these adaptive responses. Adaptive changes have been demonstrated to exist on multiple timescales and in a large variety of contexts (e.g. visual contrast adaptation, auditory stimulus specific adaptation). Here we propose that a diverse range of reported adaptive responses can be defined in terms of the accumulation of stimulus statistics by sensory filters. Using membrane potential responses to Poisson noise stimuli that varied in their stimulus statistics, we were successfully able to predict the adaptive responses to a fixed-interval stimulus across a variety of stimulus frequencies. Furthermore, we were able to achieve this across sensory cortex: in S1, V1 and A1.

Sensory adaptation has been shown to enhance change detection in behavioral tasks (Goble & Hollins, 1993; Musall et al.,2014; Tannan et al., 2007). Here, we uncovered frequency transition responses whose amplitude and length are related to the neurons’ previously adapted state. Neurons in our dataset showed a transition response amplitude in a pattern that varied between a linear or a Weber’s law like response across all modalities. These results demonstrate how adaptive processes act to preserve essential frequency information across long timescales.

There was a noticeable degree of variation both between and within modalities. For instance, A1 neurons did not display the same changes in response complexity and envelope with stimulus frequency that occur in V1 and S1 neurons. Some differences across sensory modality such as these should be expected, as each system is likely calibrated to deal with the specific statistics of its relevant sensory information. Our simple model, however, was able to capture these differences in response dynamics.

Our recordings were all performed in primary sensory areas in cortex. It is likely, however, that some of the effects described either originate or are influenced by subcortical areas, potentially as far back as the receptors themselves. Work employing similar stimuli while recording from retinal ganglion cells has found some similar phenomena to those we described, notably the termination response (Schwartz et al., 2007; Schwartz et al., 2008). This may differ by sensory system, as a recent study using fiber photometry failed to find a termination (or echo) response in the auditory thalamus (Li et al., 2017). Both the biphasic response to individual pulses (or low frequency stimulation) and response attenuation at high stimulus frequencies have also been reported subcortically (Chung et al., 2002; Funayama et al., 2016; Martin-Cortecero, J. & Nuñez, A., 2014). Another possible factor that may explain differences across modalities is the intensity of stimulation, which was not calibrated to evoke similar response magnitude across systems. Indeed, our previous studies of the somatosensory system showed that stimulus intensity entails different adaptation profiles, already observed in the trigeminal nerve (Ganmor et al, 2010, Mohar et al, 2013). Cortical mechanisms, and specifically inhibition can determine the degree of recovery from adaptation (Cohen-Kashi Malina, 2013). Inhibitory effects can vary across cortical areas and thus differently shape their adaptation behavior. Our data do not address the degree to which these response patterns may be inherited from subcortical or peripheral areas, or how this may differ by sensory modality. We instead highlight that the shared response dynamics reflect a common transformation that emerges despite the differences in transduction across sensory systems. Recent decision-making paradigms in rodents have used similar punctate stimuli in auditory, visual, and multisensory tasks, but these experiments have focused on higher-order cortical computations rather than primary sensory representations (Brunton et al., 2013; Raposo et al., 2014; Hanks et al., 2015).

A key aspect of our approach is the standardization of experimental paradigms across sensory modalities. This allows investigation to go beyond modality-specific analysis of neuronal computations to that of the underlying algorithms utilized across systems. Here we focused on adaptation, but this approach may have value in the study of other processes common in sensory cortex. For example, forms of contrast gain control, which has historically been studied in the visual cortex (Ohzawa et al., 1985), have been reported in the auditory (Cooke et al., 2018; Rabinowitz et al. 2011) and somatosensory cortices (Garcia-Lazaro et al. 2007). Understanding of this phenomenon may benefit from a cross-modal approach such as the one employed here.

The homology of cortical circuits had led many to hypothesize that different regions of cortex share a computational framework (Douglas & Martin, 2004). A certain degree of specialization among cortical areas is to be expected (Yang & Zador, 2012), but it is possible that one defining difference among cortical areas is simply the input each region receives (Sharma et al., 2000). Our results demonstrate that each of these primary sensory areas integrates information across multiple comparable timescales, yielding complex response dynamics. A change in the state of the network is therefore not required for these complex dynamics to occur, but rather can be understood as the interplay of multiple static sensory filters that span a range of relevant time scales.

## Acknowledgements

Funding was provided (to A.F., I.L., I.N. and N.P.) by the Human Frontiers Science Program and the Washington Research Foundation (UW Institute of Neuroengineering, ALF).

